# Somatostatin-Expressing Interneurons Co-Release GABA and Glutamate onto Different Postsynaptic Targets in the Striatum

**DOI:** 10.1101/566984

**Authors:** Stefano Cattaneo, Mattia Zaghi, Ripamonti Maddalena, Francesco Bedogni, Alessandro Sessa, Stefano Taverna

## Abstract

The functional contribution of somatostatin-expressing interneurons (SST-INs) to the synaptic organization of the striatum is poorly understood. Using electrophysiological recordings, optogenetic stimulation, and single-cell PCR analysis, we investigated functional patterns of synaptic connectivity in striatal SST-INs expressing channelrhodopsin-2. Photostimulation of these cells induced both glutamatergic excitatory postsynaptic currents (EPSCs) and GABAergic inhibitory postsynaptic currents (IPSCs) in striatal spiny projection neurons (SPNs) and fast-spiking interneurons (FSIs). The two synaptic components showed equally fast onset latencies, suggesting a mechanism of co-transmission. Accordingly, single-cell PCR analysis revealed that individual striatal SST-INs expressed mRNAs for both glutamate and GABA vesicular transporters (VGLUT1 and VGAT, respectively). During relatively prolonged optical stimuli (0.5-1s), IPSC arrays consistently outlasted EPSCs. As a result, photostimulation of SST-INs caused a transient burst of action potentials followed by a prolonged inhibition in postsynaptic cells.

These data suggest that striatal SST-INs are specialized to locally project synapses exerting a composite excitatory and inhibitory effect through GABA/glutamate co-transmission onto different postsynaptic targets.

## Introduction

Somatostatin-expressing interneurons (SST-INs) are known as a class of GABAergic interneurons residing in several areas of the brain (Kawaguchi et al., 1995; Freund and Buzsaki, 1996; Katona et al., 1999; Wang et al., 2004; Somogyi and Klausberger, 2005; Rudy et al., 2011; Sturgill and Isaacson, 2015; Urban-Ciecko and Barth, 2016; Ramaswamy et al., 2017). They mostly provide dendritic inhibition (Freund and Buzsaki, 1996; Pouille and Scanziani, 2004; Lovett-Barron et al., 2012; Royer et al., 2012), play an essential role in sensorimotor integration, learning, and memory (Gentet et al., 2012; Lovett-Barron et al., 2014; Stefanelli et al., 2016), and are involved in a variety of neurological disorders (Schmid et al., 2016; Zhang et al., 2016; Fuchs et al., 2017). SST-INs also represent an important subtype of interneurons in the striatum, the main input area of the basal ganglia (BG) which plays a critical role in reinforcement learning, voluntary movement, goal-directed behavior, and development of habits (Yin et al., 2005; Balleine et al., 2007; Surmeier et al., 2009; Gremel and Costa, 2013). Striatal SST-INs display neurochemical and electrophysiological properties, such as high input resistance and Ca^2+^ channel-mediated low-threshold spiking, or LTS (Kawaguchi, 1993; Partridge et al., 2009; Tepper et al., 2010; Beatty et al., 2012; Elghaba et al., 2016; Holley et al., 2019), broadly similar to those described in cortical and hippocampal SST-INs (Oliva et al., 2000; Taverna et al., 2005; Ma et al., 2006; Xu et al., 2006; Yekhlef et al., 2015; Nigro et al., 2018), and distinct from those of other major cell types such as spiny projection neurons (SPNs, (Taverna et al., 2008; Kreitzer, 2009), fast-spiking interneurons (FSIs; (Koos and Tepper, 1999; Russo et al., 2013), and other striatal populations (Tepper et al., 2018). SST-INs are generally known as inhibitory cells using GABA as a primary neurotransmitter and co-expressing several neuromodulators including somatostatin, neuropeptide-Y (NPY) and nitric oxide (NO) (Cannizzaro et al., 2003; Galarraga et al., 2007; Lopez-Huerta et al., 2008; Blomeley et al., 2015; Rafalovich et al., 2015). Anatomically, these neurons display large, sparsely branched dendrites and a very stretched axonal arborization (Kawaguchi, 1993; Ibanez-Sandoval et al., 2011). These properties suggest that SST-INs make synaptic contacts with cellular targets which may reside relatively far away from the presynaptic soma, posing a challenge to the investigation of synaptic properties with paired recordings. Recent studies have obviated this problem using optogenetic stimulation, reporting new insights in the synaptic organization of these cells and stressing their preference for long-distanced postsynaptic targets (Straub et al., 2016). However, the influence of SST-INs on postsynaptic firing activity has not been described in detail yet, leaving a gap in our understanding of striatal circuits. Here we investigated functional synaptic release properties of intrastriatal SST-INs in mouse brain slices using patch clamp recordings, optogenetic stimulation, and single-cell PCR analysis. We found that optical stimulation of these cells resulted in an efficient co-transmission of GABA and glutamate, evoking simultaneous arrays of EPSCs and IPSCs and generating a consistent firing activation-inhibition sequence in postsynaptic SPNs and FSIs. These properties were more evident in relatively older mice (>p45) and were not shared with cortical and hippocampal SST-INs, which were purely inhibitory.

The atypical, fast excitatory-inhibitory sequence induced by GABA/glutamate co-transmission following SST-IN activation adds an important contribution to the complexity of synaptic integrative properties in striatal principal cells and fast-spiking interneurons.

## Materials and Methods

### Slice preparation and electrophysiology

Recordings were performed in cortico-striatal slices prepared from a recombinant Cre-lox mouse line obtained by crossing B6.Cg-Gt(ROSA)26Sortm27.1(CAG-COP4*H134R/tdTomato)Hze/J mice (Ai27D, JAX stock number: 012567; Jackson Laboratory, Bar Harbor, ME, USA) with B6N.Cg-Ssttm2.1(cre)Zjh/J (Sst-IRES-Cre, JAX stock No. 018973). For FSI photostimulation we crossed Ai27D mice with 129P2-Pvalbtm1(cre)Arbr mice (PV^cre^, JAX stock No. 008069). The offsprings, which appeared viable and healthy, selectively expressed channelrhodopsin-2 (ChR2) in SST-or parvalbumin (PV)-expressing cells, respectively. All procedures were approved by the Italian Ministry of Health and the San Raffaele Scientific Institute Animal Care and Use Committee in accordance with the relevant guidelines and regulations. Mice of both sexes (30-90 days of age) were anesthetized with an intraperitoneal injection of a mixture of ketamine/xylazine (100 mg/kg and 10 mg/kg, respectively) and perfused transcardially with ice-cold artificial cerebrospinal fluid (ACSF) containing (in mM): 125 NaCl, 2.5 KCl, 1.25 NaH_2_PO_4_, 2 CaCl_2_, 25 NaHCO_3_, 1 MgCl_2_, and 11 D-glucose, saturated with 95% O_2_ and 5% CO_2_ (pH 7.3). After decapitation, brains were removed from the skull and 300μm-thick parasagittal slices (horizontal for thalamic stimulation experiments) were cut in ACSF at 4°C using a VT1000S vibratome (Leica Microsystems, Wetzlar, Germany). Individual slices were then submerged in a recording chamber mounted on the stage of an upright BX51WI microscope (Olympus, Japan) equipped with differential interference contrast optics (DIC). Slices were perfused with ACSF continuously flowing at a rate of 2-3 ml/min at 32°C. Whole-cell patch-clamp recordings were performed in dorsolateral striatum and somatosensory cortex using pipettes filled with a solution containing the following (in mM): 10 NaCl, 124 KH_2_PO_4_, 10 HEPES, 0.5 EGTA, 2 MgCl_2_, 2 Na_2_-ATP, 0.02 Na-GTP, (pH 7.2, adjusted with KOH; tip resistance: 4-6 MΩ). All recordings were performed using a MultiClamp 700B amplifier interfaced with a PC through a Digidata 1440A (Molecular Devices, Sunnyvale, CA, USA). The liquid junction potential was not corrected. The series resistance was partially compensated (40-50%) using the amplifier control circuit. Data were acquired using pClamp10 software (Molecular Devices) and analyzed with Origin 9.1 (Origin Lab, Northampton, MA, USA). Voltage-and current-clamp traces were sampled at a frequency of 30 kHz and low-pass filtered at 2 kHz. The cell input resistance (R_in_) was calculated by dividing the peak voltage change value in response to the injection of a hyperpolarizing current step (−100 pA) in current clamp recordings. Latencies between optical stimuli and ChR2-induced direct or synaptic currents were measured as the time gap between stimulus trigger start (recorded as a digital output trace in pClamp) and current onset, defined as the time at which membrane potential traces deflected by a value equal to at least twice the standard deviation (SD) of the baseline mean value preceding the stimulus. Compound IPSC and EPSC durations were measured as the time interval between current onset and earliest time point marking a full decay of the last PSC to baseline.

### Photostimulation of ChR2-expressing interneurons

Optical stimuli were generated using a diode-pumped solid state laser (wavelength: 473 nm; Shanghai Dream Lasers Technology, Shanghai, China) connected to the epi-illumination port of the microscope through a multi-mode optical fiber. The beam was deflected by a dichroic mirror and conveyed to the slice through a 40x water-immersion objective (spot size: 0.06 mm^2^). The light power measured with an optical power meter at the level of the slice surface was ~2 mW, yielding a light density value of ~33 mW/mm^2^. Photostimuli were TTL-triggered using Clampex digital output signals.

### Single-cell polymerase chain reaction (PCR)

Neurons in slices obtained from mice aged p55-p65 were visually identified and their firing patterns were recorded in order to confirm their functional identity. Patch pipettes were filled with an autoclaved internal solution containing the following (in mM): 140 KCl, 5 EGTA, and 5 HEPES, pH 7.3 with KOH. After recording, the cytoplasm was harvested into the pipette by gently applying negative pressure. An initial reverse transcription (RT) reaction was conducted after pressure ejection of the cytoplasm into a microcentrifuge tube containing the REPLI-g WTA single cell kit (Qiagen). Each cell was incubated in a total volume of 5.5 μl at 24°C for 5 min and then cooled to 4°C. Cells were subsequently treated for 10 min at 42°C with 1 μl gDNA wipeout buffer before adding 3.5 μl RT mix to synthesize first strand cDNA (RT mix: 0.5 μl oligodT primer, 2 μl RT buffer, 0.5 μl random primer and 0.5 μl RT enzyme mix). The tubes were incubated at 42°C for 1 h, then at 95°C for 3 min. A ligation step was subsequently carried out at 24°C for 30 min with 5 μl ligation mix (4 μl ligase buffer and 1 μl ligase mix) for each sample. The reaction was stopped by incubating at 95°C for 5 min. The final amplification step was performed at 30°C for 2 h after adding the amplification mix (15 μl buffer and 0.5 μl REPLI-g SensiPhi DNA polymerase), eventually raising the temperature to 65°C for 5 min to stop the reaction.

Single cell PCRs were performed on diluted cDNA (1:100) using Go-Taq polymerase (Promega) with the following protocol: [94° 5’ – 35x(94 30” – 60 30” – 72 1’30”) – 72° 7’] For *VGAT*, a nested PCR was performed where the final amplification was made using the protocol described above while the following protocol was used as first step: [94° 5’ – 35x(94 30” – 55 30” – 72 1’30”) – 72° 7’]. Every PCR was loaded on 2% agarose gel for visualization. *VGAT* and *VGLUT1* bands were extracted from gel and further examined by Sanger sequencing (Microsynth AG, Balgach, Switzerland).

The Following primers were used:

*Slc32a1*(*VGAT)*_1st_ F: 5’-GAGCATCGCCAGGGCCTGCA-3’ |
*Slc32a1*(*VGAT)*_1st_ R: 5’-CCGCTCACCACGACGTACAA-3’ |
*Slc32a1*(*VGAT)*_2nd_ F: 5’-CGACAAACCCAAGATCACGG-3’ |
*Slc32a1(VGAT)* _2nd_ R*: 5’-* AGGATGGCGTAGGGTAGGC-3’ |
*Sst_*F: 5’-ACCGGGAAACAGGAACTGG-3’ |
*Sst_R*: 5’-TTGCTGGGTTCGAGTTGGC-3’ |
*Slc17a7(VGLUT1)_*F: 5’-CAGGAGGAGTTTCGGAAGCTGG-3’ |
*Slc17a7(VGLUT1)_*R: 5’-GCGATGATGTAGCGACGAG-3’ |
*Slc17a6(VGLUT2)_*F: 5’-TGGAAAATCCCTCGGACAGAT-3’ |
*Slc17a6(VGLUT2)_*R: 5’-CATAGCGGAGCCTTCTTCTCA-3’ |
*Slc17a8(VGLUT3)_*F: 5’-TTCCCGGTGGCTTCATTTCAA-3’ |
*Slc17a8(VGLUT3)_*R: 5’-ACAGCCGTAATGCACCCTC-3’ |
*Eomes_*F: 5’-TCCAGGGGGGCAAGTGGGTG-3’ |
*Eomes_*R*: 5’-*CATCTGTGTGTTGTTGTTTG-3’ |
*18S_*F: 5’-GGTGAAATTCTTGGACCGGC-3’ |
*18S_*R: 5’-GACTTTGGTTTCCCGGAAGC-3’ |

### Immunohistochemistry

Mice were intracardially perfused using a peristaltic pump connected to a needle inserted in the left ventricle. We first injected 10 ml of cold ACSF followed by 20 ml of paraformaldehyde (PFA) dissolved at 4% w/v in phosphate-buffered saline (PBS) at 4°C. Brains were then quickly removed and post-fixed in PFA overnight at 4°C. Sagittal sections (15μm thick) were cut using a cryostat (Leica). Slices were permeabilized with a blocking solution containing 0.3% Triton-X100, 1% bovine serum albumin, or BSA, and PBS, and incubated with primary antibodies diluted in blocking solution (anti-SST 1:100 v/v, Millipore, and anti-red fluorescent protein 1:400 v/v, Rockland Immunochemicals, overnight at 4°C). Slices were then washed for 1h by alternating PBS and blocking solution every 15 min, and subsequently incubated with secondary antibodies diluted 1:200 v/v in blocking buffer solution for 2h at room temperature (anti-rat ALexa488, anti-mouse Alexa546, Jackson Immunoresearch). After incubation with secondary antibodies, slices were washed for 2h by alternating blocking solution and PBS every 15 min, then rinsed in PBS and mounted with an anti-fading agent (Fluor Safe, Millipore). Images were collected using a Leica SP5 confocal microscope.

### Statistical analysis

Results were compared using paired or unpaired Student’s t-tests and ANOVA for normally distributed datasets, and Mann-Whitney rank sum tests otherwise (SigmaStat, Systat Software, Chicago, IL). Population rates were compared using a Z-score test for two populations. Results are given as means ± s.e.m. in text and represented by box plots in figures. Boxes include 25^th^ and 75^th^ percentiles (horizontal edges), median value (inner line), and min-max values (whiskers). Differences were considered significant at p<0.05.

### Drugs

All drugs were obtained from Sigma except 2,3-dioxo-6-nitro-1,2,3,4-tetrahydrobenzo[*f*]quinoxaline-7-sulfonamide disodium salt (NBQX) and 2-(3-carboxypropyl)-3-amino-6-(4 methoxyphenyl)pyridazinium bromide (gabazine) which were obtained from Hello Bio (Bristol, UK).

## Results

### Photostimulation specifically activates SST-INs

Individual striatal cells were first recorded in order to assess the typical identity of SST-INs (Kawaguchi et al., 1995; Kubota and Kawaguchi, 2000; Beatty et al., 2012) in fluorescent cells in slices prepared from SST-ChR2 mice. Visually identified, red fluorescent cells invariably displayed the electrical properties of SST-INs, namely a relatively high R_in_ (350 ± 31 MΩ, n = 30), a prominent depolarizing sag in response to injection of hyperpolarizing current pulses, and post-inhibition rebound firing (Figure 1A). The great majority of these cells (29 out of 30, 97%) readily fired trains of action potentials in response to direct illumination with blue light. We found some variability in the frequency response of SST-INs to either current injection or photostimulation, so that a fraction of cells (19 out of 30, 63%) displayed a relatively regular AP firing pattern with little or no adaptation, while the remainder (11 out of 30, 37%) were more irregular or strongly adapting. In any case, ChR2 expression was highly co-localized (>80%) with anti-SST immunohistochemical labeling (Figure 1B; (Yekhlef et al., 2015)). Conversely, all recorded cells with electrophysiological features typical of SPNs (N = 130; (Taverna et al., 2008)), FSIs (N = 26; (Russo et al., 2013)), cholinergic interneurons (N = 10; (Wilson et al., 1990)), and cortical principal cells (N = 12; (Yekhlef et al., 2015)) displayed no fluorescence associated with ChR2 expression and lack of light-induced direct depolarization, suggesting that photostimulation robustly activated SST-INs with a high degree of specificity.

**Figure 1.**
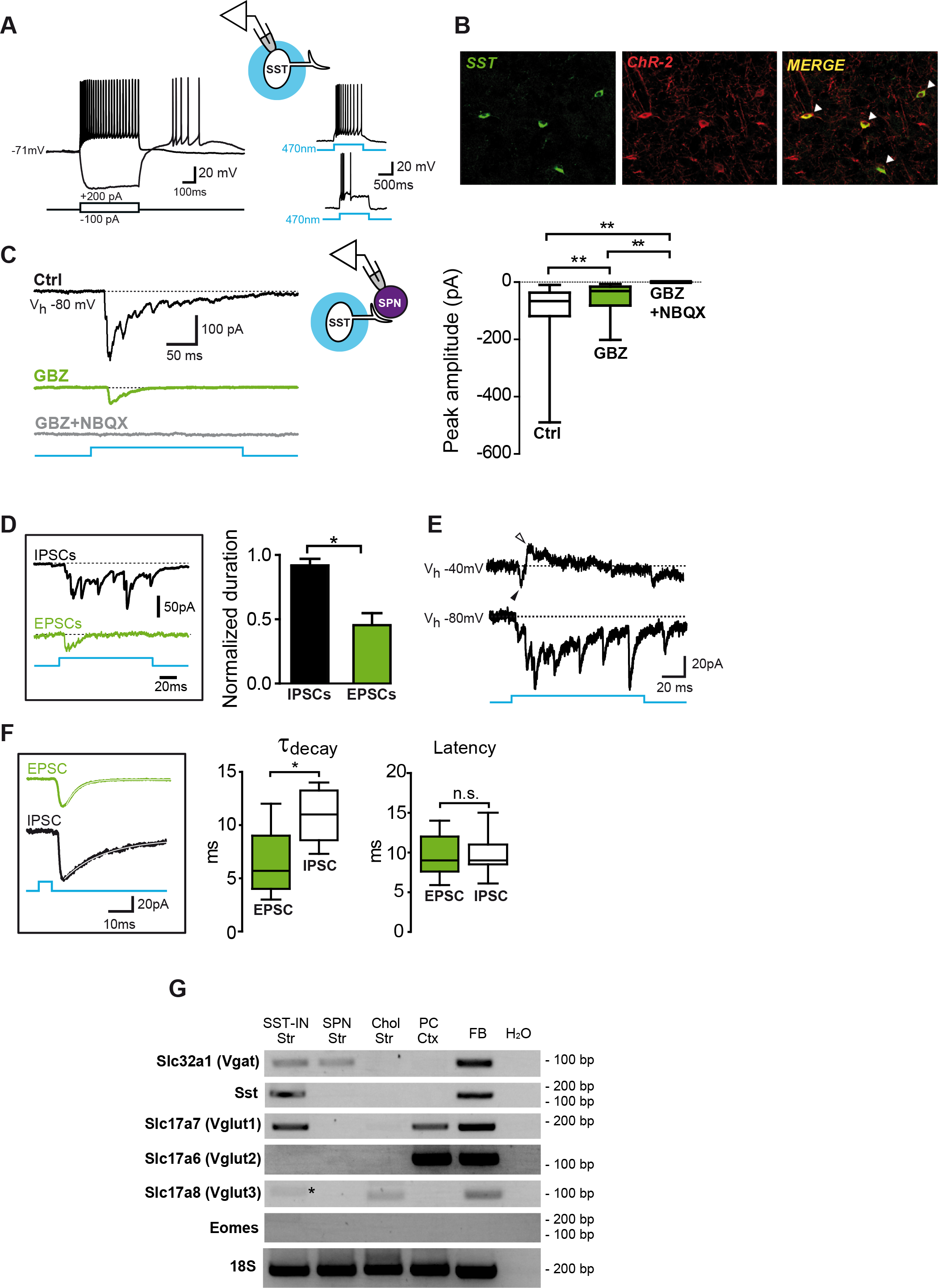
GABAergic and glutamatergic postsynaptic currents (PSCs) evoked by SST-IN photostimulation. **A:** electrical properties of a regular-spiking SST-INs in response to injection of current steps (500 ms, left) and direct photostimulation with blue light at 470nm (1s, right). The upper trace on the right is from the same cell as that in the left panel, while the bottom is from a different SST-INs showing an irregular firing pattern in response to light. **B:** confocal microphotographs showing examples of anti-SST immunostaining co-localizing with ChR-2 expression. **C:** voltage clamp traces representing a compound PSC (Ctrl) recorded in a striatal SPN in response to SST-IN photostimulation. Extracellular perfusion with the GABA_A_-receptor antagonist gabazine (GBZ, 10µM) unmasked a smaller, rapidly-depressing compound EPSC that was completely blocked by the AMPA-receptor antagonist NBQX (5µM). Summary plots of PSC peak amplitudes are shown on the right (**p<0.01). **D:** pharmacologically isolated compound IPSCs and EPSCs evoked by a 200-ms-lasting flash. The histogram shows a significantly shorter duration for compound EPSCs (normalized to flash duration) as compared to IPSCs. **E**: Examples of light-induced post-synaptic currents recorded at two different holding potentials (V_h_) in plain ACSF. At V_h_ = −40mV the GABAergic component of the PSC was visible as an outward current (white arrowhead). The black arrowhead marks an inward EPSC. **F**: individual EPSC and IPSC recorded in response to brief flashes (5ms) in the presence of GBZ and NBQX, respectively (V_h_ = −80 mV). The box plots summarize decay time constants and latencies for individual EPSCs and IPSCs. **G**: representative gel of single-cell RT-PCR analysis showing mRNA expression of *Slc32a1, Sst, Slc17a7, Slc17a6, Slc17a8, Eomes* (an embryonic gene not expressed in the adult, used as a negative control) and *18S* (a housekeeping gene as positive control). An expression pattern example for each of the following cell types is shown: striatal somatostatin-expressing interneuron (SST-IN Str), striatal spiny projection neuron (SPN Str), striatal cholinergic interneuron (Chol Str), cortical principal cell (PC Ctx). PCR performed on mRNA extract from adult forebrain tissue (FB) and blank reaction (H_2_O) are also shown as positive and negative controls, respectively. The asterisk indicates a non-specific PCR band.

### SST-INs co-release both GABA and glutamate onto SPNs

Photostimulation of SST-INs elicited a composite synaptic response in individual postsynaptic cells. During voltage-clamp recordings in SPNs at a V_h_ of −80 mV, blue light flashes (0.5-1s) evoked a compound inward current (mean peak amplitude: −113 ± 16 pA, n = 53; Figure 1C, black trace). After extracellular perfusion with the GABA_A_ receptor antagonist gabazine (GBZ, 10 µM), a smaller but consistent residual compound current was unmasked (mean peak amplitude: −59 ± 13 pA; p<0.01, n = 30, Mann-Whitney Rank Sum Test; Figure 1C, green trace; we used an unpaired statistical test since in some cases cells were recorded when GBZ was already present in the extracellular ACSF). Such current was completely inhibited by the AMPA-receptor antagonist NBQX (5µM; Figure 1C, gray trace), suggesting that SST-IN photostimulation induced synaptic release of both glutamate and GABA that were functionally active on SPN postsynaptic receptors.

Interestingly, the average duration of the AMPA-receptor-mediated compound current (I_AMPA_ –normalized to the flash duration, since in some cases we used different stimulation lengths across experiments) was significantly shorter than the GABA_A_-R-mediated component (I_GABA_) recorded in 5µM NBQX (40 ± 4% *vs*. 94 ± 4%, respectively, n = 10, p = 0.001, unpaired t test; Figure 1D). I_AMPA_ and I_GABA_ were also distinguished by their different reversal potentials (E_rev_), as both inward EPSCs (estimated E_rev_= 0 mV) and outward IPSCs (estimated E_rev_= −60mV) were detected during relatively prolonged flashes (150 ms) at an intermediate V_h_ (−40 mV; Figure 1E). In addition, individual EPSCs elicited with brief light pulses (5 ms) at V_h_ = −80 mV in the presence of GBZ had a significantly faster decay time constant (τ_dec_) than IPSCs recorded at the same V_h_ in the presence of NBQX (6.3 ± 1.0 ms *vs*. 10.8 ± 1.0 ms, respectively, n = 7, p<0.05, unpaired t test; Figure 1F, *inset*), confirming the identity of the two synaptic components. Importantly, the average latencies of EPSCs and IPSCs were nearly identical (9.4 ± 0.6 ms vs. 9.7 ± 1.0 ms, respectively, n = 7, p=0.8, unpaired t test; Figure 1F). Altogether, these data suggest that both GABA and glutamate are simultaneously released onto striatal SPNs in response to photoactivation of SST-INs. However, the glutamatergic response is relatively fast-depressing and persists for a significantly shorter time than the inhibitory component.

To confirm the presumed co-transmission of the two different neurotransmitters and identify its molecular basis, we performed semi-quantitative reverse transcription-PCR (RT-PCR) in cytoplasmic samples of identified individual cells collected through the recording pipette. SST-INs consistently expressed *Sst* (the gene encoding for somatostatin; 10 out of 10 cells; Figure 1G). In a subset of SST-INs (3 out of 10 cells) we detected co-expression of both GABA and glutamate vesicular transporters, VGAT (*Slc32a1*) and VGLUT1 (*Slc17a7)*, respectively (Figure 1G, Supplementary Figure S1). Conversely, striatal SPNs expressed VGAT but not VGLUT (3 out of 3 cells), while cortical principal cells (PCs) expressed VGLUT1 and VGLUT2 (*Slc17a6*), but not VGAT (3 out of 3 cells). SST-INs neither expressed VGLUT2 (0 out 10 cells) nor VGLUT3 (*Slc17a8;* 0 out of 10 cells). One striatal cholinergic interneuron was positive for VGLUT3 (1 out of 1 cell), consistently with previous findings (Fremeau et al., 2002; Gras et al., 2002; Nelson et al., 2014).

These data suggest that, at least in a subset of striatal SST-INs, GABA/glutamate co-transmission is supported by the concomitant expression of vesicular transporters VGAT and VGLUT1, respectively.

### GABA/glutamate co-transmission is age-dependent

The probability to detect SST-IN-mediated release of glutamate was much higher in relatively older than younger mice. We recorded compound EPSCs (in the presence of GBZ) in 18 out of 20 SPNs (90%) from 8 mice aged 45 to 90 days. Conversely, in 7 younger animals (p30-p44) EPSCs were only detected in 15 out of 27 SPNs (56%, p=0.028, Z-test for two population proportions; Figure 2A). In addition, the median EPSC amplitude in responding SPNs was significantly larger in older than younger mice (−81 pA *vs.* −17 pA, respectively, n = 18 and 15, p=0.003, Mann-Whitney Rank Sum Test; Figure 2A).

**Figure 2.**
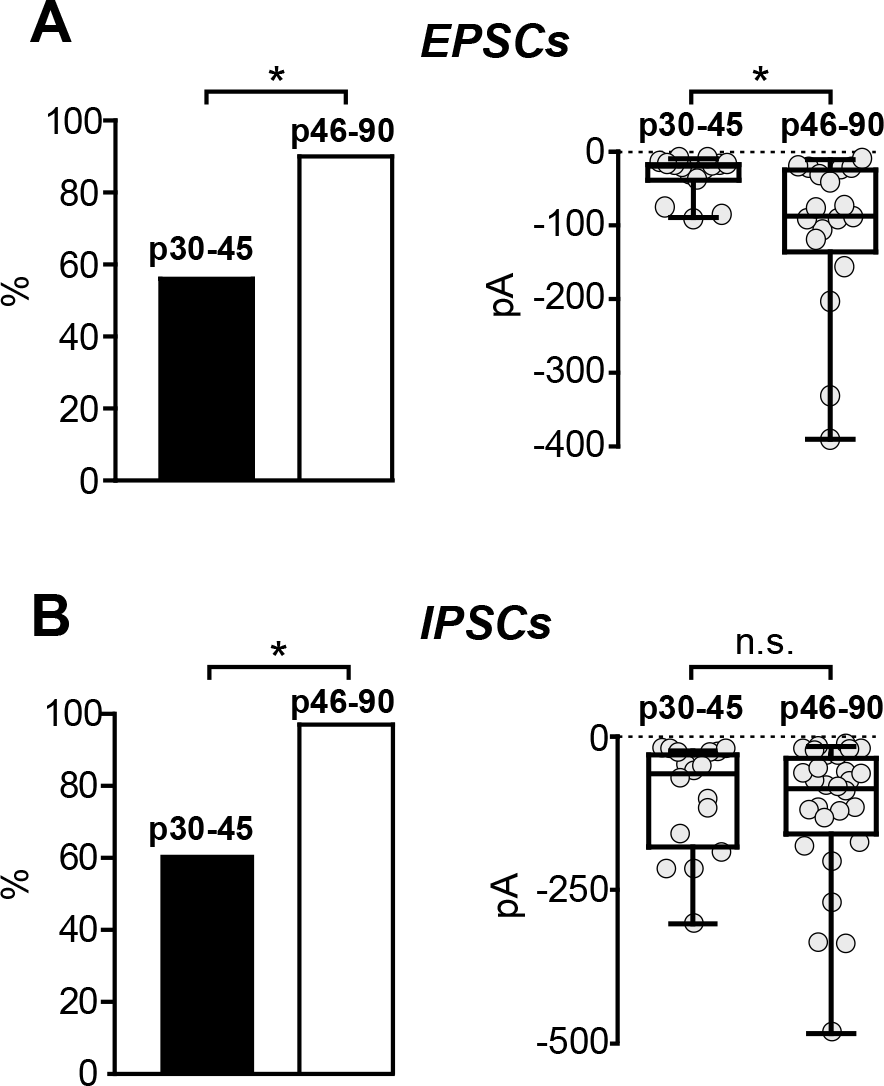
The expression of both glutamatergic and GABAergic PSCs in response to SST-IN photostimulation is age-dependent. **A**: left, rates of successfully detected compound EPSCs (recorded in the presence of 10µM GBZ) in striatal SPNs from animals grouped in two age cohorts (postnatal day 30 (p30) through p45, and p46 through p90). Right, summary box plots of compound EPSC peak amplitudes. Each dot represents the amplitude of a compound EPSC recorded in one individual SPN. **B**: same as in A, but for compound IPSCs recorded in the presence of 5µM NBQX (n.s.: not significant).

Similarly, compound IPSCs (in the presence of NBQX) were found in 30 out of 31 SPNs (97%) from 8 older mice (p45-p90) and in 17 out of 28 SPNs (61%) from 3 younger mice (p30-p44; p=0.002, Z-test for two population proportions; Figure 2B). The median IPSC amplitude was also larger in the older group, though the difference was not statistically significant (−85 pA *vs.* −60 pA, respectively, n = 30 and 18, p=0.57, Mann-Whitney Rank Sum Test; Figure 2B).

Thus, both glutamate and GABA release by SST-INs become progressively more robust well into the adult age in mice.

### SST-IN photoactivation promotes a transient excitation followed by robust inhibition of SPN firing activity

The SST-IN-mediated dual synaptic response described above might unconventionally affect the firing activity of postsynaptic SPNs. To assess this, we recorded trains of action potentials induced by steady, suprathreshold DC injection (200-300 pA) in individual SPNs in current-clamp configuration. Once a stable firing activity was reached, we delivered blue light flashes (2-4 s) which consistently caused a transient increase in AP frequency lasting 100-300ms, followed by a more prolonged inhibition that persisted up to the flash offset (Figure 3A). The cell would then gradually recover to its baseline frequency. Mean frequency values measured during light delivery were significantly different from pre-and post-flash ones (pre-flash: 6 ± 0.8 Hz, early flash increase: 17 ± 4 Hz, late flash decrease: 2 ± 1 Hz, post-flash: 6 ± 1 Hz, n = 7, p<0.05, one-way ANOVA; Figure 3B). Similar results were obtained when flashes were delivered over a higher baseline frequency induced by larger DC injection (300-400 pA; ctrl: 18 ± 5 Hz, early flash increase: 44 ± 9 Hz, late flash decrease: 11 ± 5 Hz, post-flash: 16 ± 4 Hz, n = 5, p<0.05, one-way ANOVA; Figure 3B). These results suggest that co-transmission of glutamate and GABA during SST-IN stimulation induce a fast, transient excitatory effect followed by a slower inhibition of firing activity in striatal postsynaptic SPNs.

**Figure 3.**
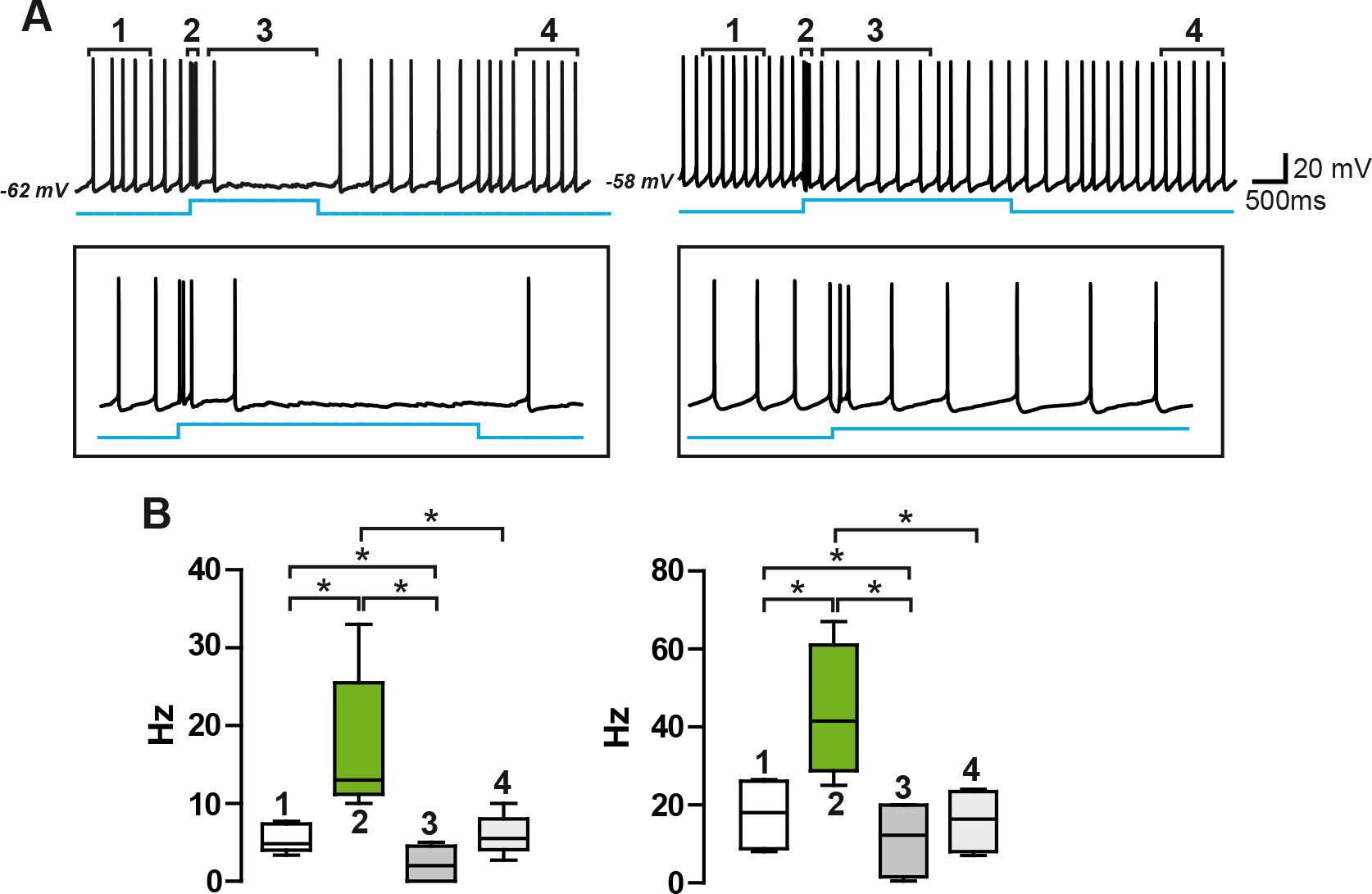
SST-IN photoactivation induces an excitation-inhibition sequence affecting SPN firing activity. **A:** examples of current clamp recordings of SPN firing activity. Steady positive DC (200-300 pA) was injected to induce action potentials at different frequencies (left and right panels, respectively). In both cases SST-IN photoactivation induced a transient increase in firing frequency followed by a pronounced inhibition. **B:** summary box plots with frequency variations induced by SST-IN photoactivation. Each box is marked by a number corresponding to trace portions indicated in A.

### GABA/glutamate co-transmission also occurs in striatal SST-IN synapses targeting fast-spiking interneurons

A similar pattern of synaptic response was detected in fast-spiking interneurons (FSIs, another major class of striatal cells) in response to SST-IN photostimulation (Figure 4). Control compound currents induced by blue light flashes (0.5-1s) were only partially reduced by 10µM GBZ (ctrl: −220 ± 84 pA, GBZ: −78 ± 19 pA, respectively, n = 6, p = 0.11, paired t test), the residual current being completely blocked by 5µM NBQX. Individual EPSCs had faster kinetics than IPSCs (τ_dec_: 1.5 ± 0.2 ms *vs*. 6.4 ± 0.7 ms, respectively, n = 6, p<0.05, unpaired t test; Figure 4B), while their latencies from pulse onset were comparable (9.0 ± 1.9 ms *vs*. 8.3 ± 0.9 ms, n = 6, p>0.05, unpaired t test; Figure 4B). Similarly to what we observed in SPNs, photoactivation of SST-INs induced a transient increase in FSI AP frequency followed by a slowdown that often persisted throughout the pulse delivery (pre-flash: 56 ± 14 Hz, early increase: 94 ± 14 Hz, late decrease: 32 ± 10 Hz, post-flash: 42 ± 11 Hz, n = 4, p<0.05, one-way ANOVA; Figure 4C). Thus, striatal SST-INs also project GABA/glutamate co-releasing synapses onto FSIs, exerting a fast excitatory-inhibitory sequence similar to that observed in postsynaptic SPNs.

**Figure 4.**
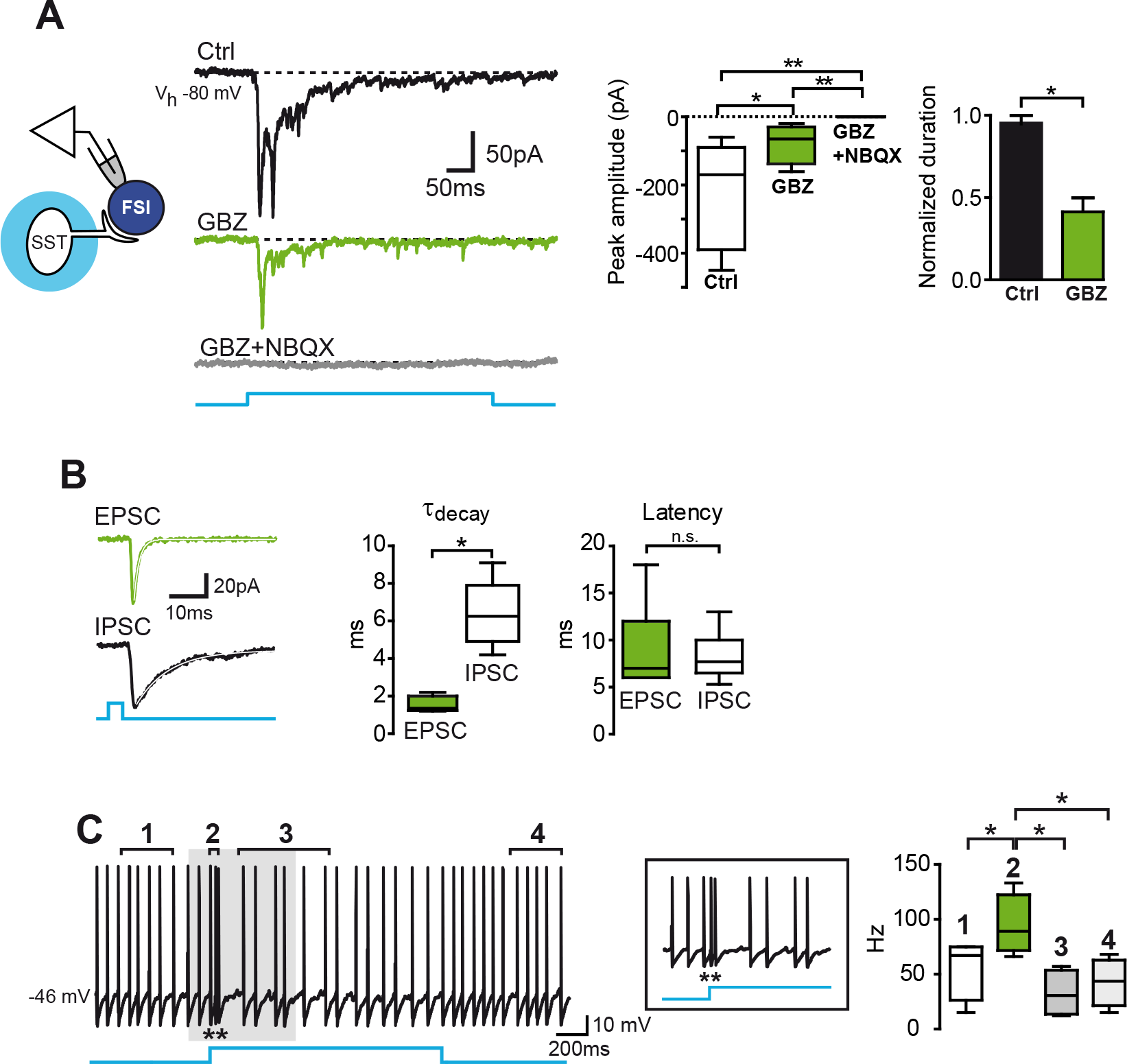
SST-INs co-release GABA and glutamate onto FSIs. **A:** voltage clamp traces showing a compound PSC recorded in a striatal FSI in response to photostimulation of SST-INs. After extracellular perfusion with 10µM GBZ a sizeable, faster inward current was still detected (green trace). No residual current was visible after the addition of NBQX (grey). Summaries of peak amplitude and normalized duration of compound currents are shown on the right. **B:** individual EPSC and IPSC with relative box plots for decay time constants and latencies. **C:** excerpt of a current clamp recording of the firing activity in the same FSI as in B. Similarly to what observed in SPNs, SST-IN photoactivation induced a transient increase in firing frequency followed by a marked slowdown. Upon light offset, the frequency was restored to a value similar to that prior to light onset.

### Presynaptic FSIs only release GABA onto SPNs

We asked whether striatal FSIs may be able to co-release glutamate and GABA in a similar manner as that seen for presynaptic SST-INs. Using slices obtained from transgenic mice expressing ChR2 in FSIs, we delivered blue light flashes over the striatum in order to specifically excite FSIs while patching postsynaptic SPNs in voltage-clamp mode at V_h_ = −80 mV (Figure 5). Differently from SST-INs, FSIs exclusively elicited GABAergic IPSCs as perfusion with 10µM GBZ left no detectable current (peak amplitude, ctrl: −466 ± 145 pA, GBZ: −2 ± 2 pA, n = 6, p<0.05, paired t test, Figure 5A). Accordingly, during FSI photoactivation we observed a purely inhibitory effect on SPN firing activity, with no sign of the transient excitation that was consistently recorded during SST-IN stimulation (pre-flash: 8.7 ± 1.6 Hz, during flash: 1.7 ± 1.1 Hz, post-flash: 7.0 ± 1.6 Hz, n = 6, p<0.05, one-way ANOVA; Figure 5B). Thus, of these two major classes of striatal GABAergic interneurons only SST-INs are capable to co-release glutamate and GABA onto SPNs.

**Figure 5.**
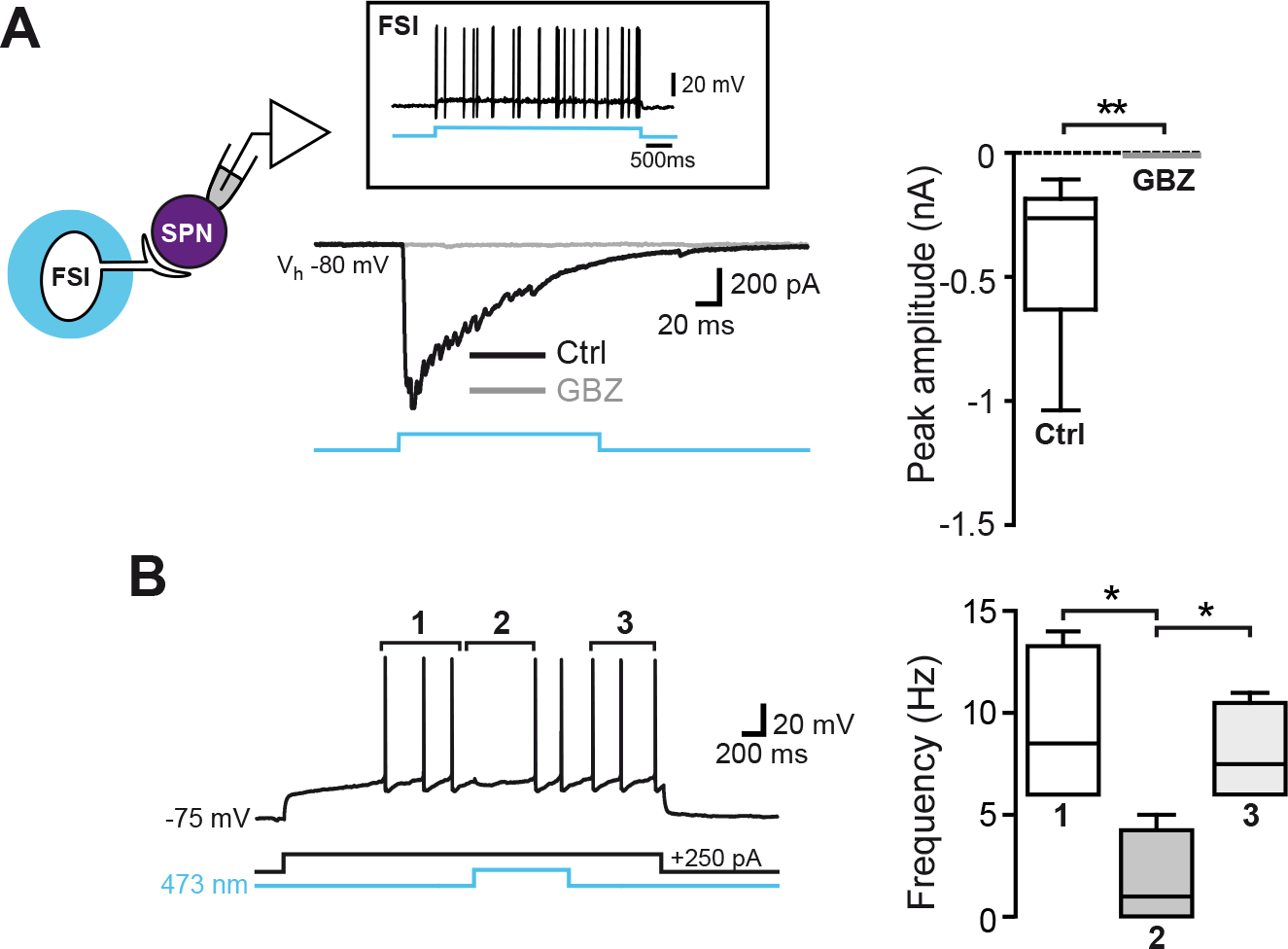
FSIs release GABA, but not glutamate, onto SPNs. **A:** voltage clamp traces representing compound PSCs recorded in a striatal SPN in response to local FSI photostimulation. The inset on top shows an example trace recorded in a FSI firing a train of APs in response to blue light stimulation. Differently from the case of presynaptic SST-INs, extracellular perfusion with 10µM GBZ was sufficient to completely block the compound PSC. Summary box plots with median peak currents before and after GBZ are shown on the right. **B:** current clamp recording of firing activity induced by injection of a step current (250 pA, 3s) in the same SPN as in A. Spikes were transiently blocked during FSI photoactivation, suggesting a purely inhibitory role of FSI GABAergic synapses with no evidence for excitation mediated by glutamate co-release. The box plot on the right summarizes the effect of FSI activation on SPN firing frequency.

### Cortical SST-INs are exclusively GABAergic

To assess whether striatal SST-INs share their ability of co-releasing GABA and glutamate with their analog cells residing in cortical areas, we recorded synaptic currents in cortical principal cells (PCs) in response to local photostimulation of SST-INs (Figure 6). In this configuration, only GABAergic, GBZ-sensitive currents were detected (ctrl: −54 ± 13 pA, GBZ: 0 ± 0 pA, n = 5, p<0.05, paired t test; Figure 6A,B). Consequently, a strong inhibition of firing activity was the only evident effect in current-clamp recordings (pre-flash: 4.5 ± 1.0 Hz, during flash: 0.9 ± 0.3 Hz, n = 5, p<0.05, paired t test; Figure 6C,D). Similar results were obtained when recording CA1 neurons during photostimulation of hippocampal oriens-lacunosum moleculare (O-LM) SST-INs (not shown).

**Figure 6.**
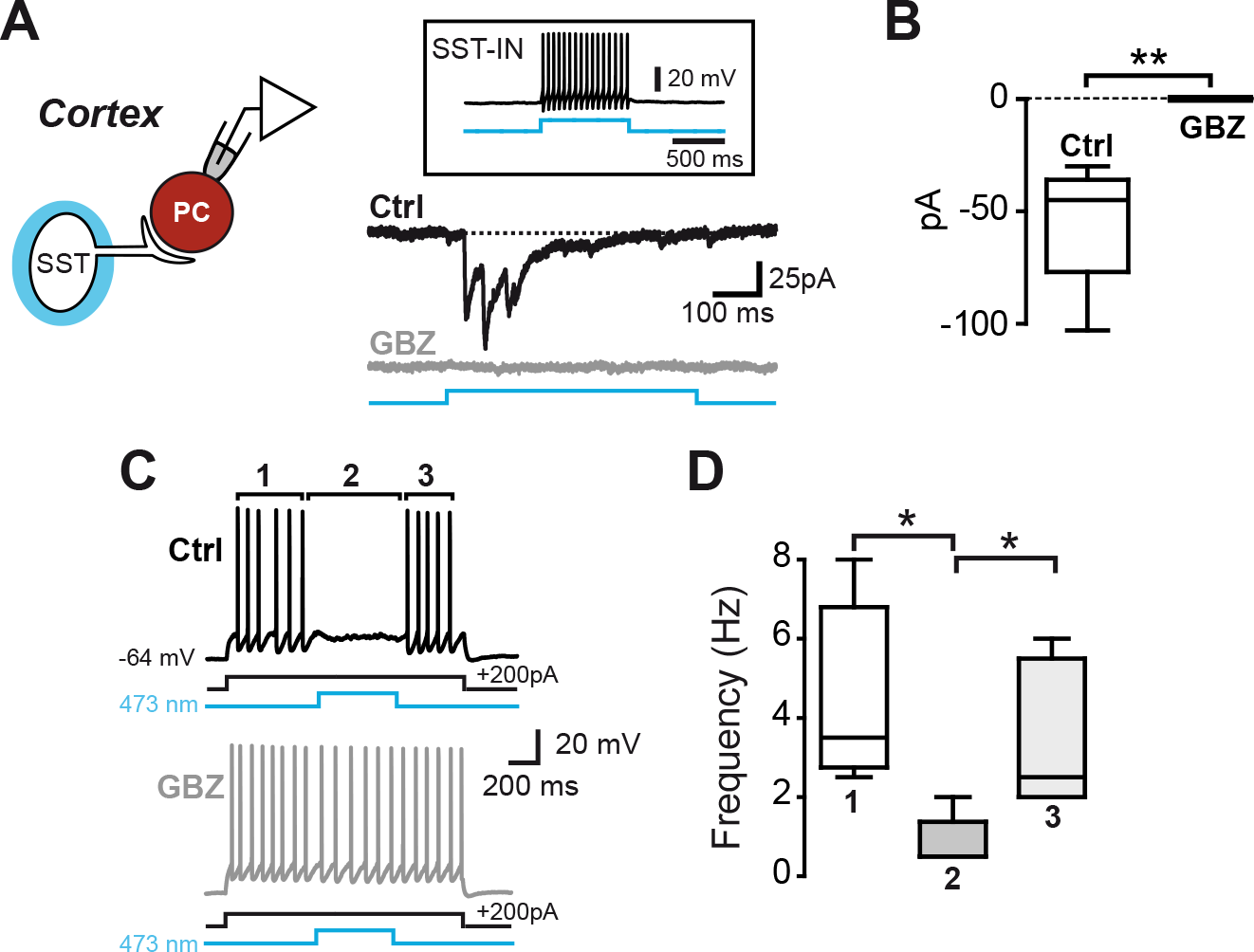
Cortical SST-INs are exclusively GABAergic. **A**,**B:** photostimulation of cortical SST-INs elicited exclusively GABAergic compound ISPCs in postsynaptic principal cells (PCs). No response to flashes was detectable after perfusion with 10µM GBZ. The inset on top shows an example trace recorded in a cortical SST-IN firing a train of APs in response to blue light stimulation. **C:** top, firing activity evoked by injection of a current step (+200 pA, 500ms) in a cortical PC was robustly blocked by photostimulation of SST-INs. Bottom, in the same cell no effect of photostimulation was detected in the presence of 10µM GBZ. **D:** summary box plot of median firing frequencies before, during, and after blue light flashes in control conditions.

### GABA/glutamate co-transmission is not mediated by synaptic projections from extra-striatal areas

Finally, we found no synaptic response (either GABA-or glutamatergic) in striatal SPNs during flash stimulation of SST-INs in various cortical, pallidal, or thalamic regions (Figure 7). We did not detect the recently reported IPSCs mediated by long-range cortico-striatal SST-INs (Melzer et al., 2017) likely because these projections preferentially target the ventro-lateral striatum, while we concentrated our recordings in the dorsal aspects of the region. We occasionally found IPSCs evoked by photostimulation of lateral cortical areas (e.g. auditory; not shown), though these responses were only GABAergic, consistently with a study by (Rock et al., 2016). Thus, our data suggest that (1) only intrastriatal SST-INs are able to co-release GABA and glutamate onto local postsynaptic targets, while their cortical and hippocampal counterparts are exclusively GABAergic, and (2) GABA/glutamate co-transmission is likely not exerted by synaptic terminals originating from extra-striatal cells located in striatally-projecting regions.

**Figure 7.**
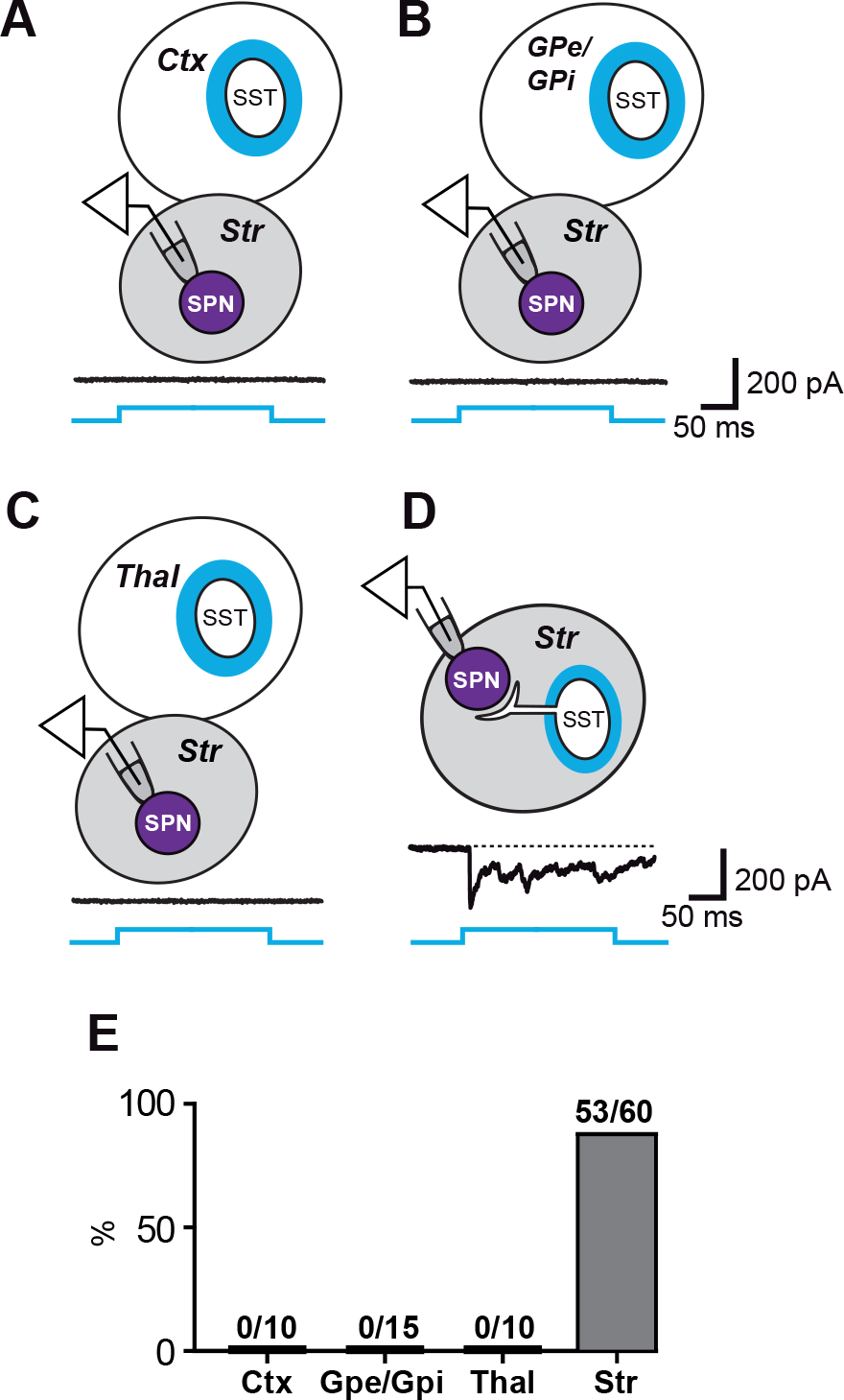
SST-IN-mediated PSCs are only detectable with intrastriatal photostimulation. **A**-**D:** light flashes delivered in motor or somatosensory cortex (Ctx), external and internal globus pallidus (GPe/GPi), or thalamus (Thal) evoked no detectable synaptic response in striatal SPNs. Conversely, intrastriatal (Str) photostimulation invariably elicited PSCs (note that any type of response was counted, including IPSCs or EPSCs only, or mixed PSCs). **E**: summary of percentages of successful PSCs obtained during photostimulation targeting any of the four different areas.

## Discussion

We report a mechanism of simultaneous co-transmission of GABA and glutamate exerted by SST-INs in the striatum–but not in cortex or hippocampus–targeting postsynaptic SPNs and FSIs. The different dynamic properties of action of the two neurotransmitters resulted in a consistent pattern of postsynaptic electrical activity perturbation, i.e. a transient excitation followed by a prolonged inhibition of action potential firing. Co-transmission of glutamate and GABA was supported by concomitant expression of vesicular transporters VGLUT1 and VGAT in at least a subset of individual SST-INs. Finally, GABA/glutamate co-transmission was consistently detected in relatively older mice (aged over 45 days), but less so in younger animals.

### Optical activation of SST-INs is reliable and specific

In a first set of experiments we assessed the specificity of ChR2 expression in our mice, in view of recent concerns regarding the use of transgenic lines resulting from crossbreeding a “floxed” cre reporter line (Ai14, JAX 007908) and an SST-IRES-cre line (JAX 013044) (Hu et al., 2013; Straub et al., 2016). In these studies, a reporter protein was found expressed in sizeable fractions of non-SST-INs, raising questions about the experimental reliability of the SST-IRES-cre mice. Here, we used a variant line (JAX 018973) re-derived on a different strain (C57BL/6NJ) from the parental one described in the studies mentioned above (C57BL/6;129S4). We found no evidence for off-target ChR2 expression as none of the cells (including cortical PCs) displaying firing properties different from those typical of SST-INs directly responded to light. In addition, we detected no synaptic currents in striatal cells in response to photostimulation of a variety of extra-striatal areas (Figure 7). Thus, in these mice optically-activated GABA and glutamate co-release seems to be exclusively mediated by striatal SST-INs.

### SST-IN selectively co-release GABA and glutamate

An increasing number of studies have recently shown that individual neurons are able to release both GABA and glutamate (Gutierrez, 2005; Root et al., 2014; Fattorini et al., 2015; Ntamati and Luscher, 2016; Tritsch et al., 2016; Granger et al., 2017; Hashimotodani et al., 2018; Root et al., 2018). Interestingly, synaptic terminals of pallidal cells projecting to the lateral habenula (LHb) were recently found to co-release the two neurotransmitters (Shabel et al., 2014; Meye et al., 2016). These terminals belong to pallidal SST-INs expressing VGLUT2 and VGAT (Wallace et al., 2017). There is an interesting analogy between these findings and our results in that SST-INs (but not other cells in the respective areas) of these two major regions of the basal ganglia (striatum and globus pallidus pars interna–or entopeduncular nucleus in rodents) appear to be a class of interneurons specialized in GABA/glutamate co-release. On the other hand, striatal SST-INs appear to project their co-releasing synapses locally, while pallidal ones project out to LHb. In addition, glutamate release is associated with VGLUT2 expression in pallidal SST-INs (Wallace et al., 2017) and VGLUT1 in striatal ones (our present results), suggesting a different molecular signature for their co-transmission ability and/or heterogeneity in their embryonic origin.

### Do all SST-INs co-release GABA and glutamate?

Our optogenetic experiments revealed GABA/glutamate co-transmission in response to the majority of successful stimulations, probably because we delivered light flashes over a relatively broad area to recruit as many SST-INs as possible. To assess the rate of individual co-releasing cells we tried to record cell pairs made of one SST-IN and one SPN (or FSI), but failed to find any functional connectivity at all in 18 different pairs –likely due to the relatively long distance between connected pairs of this kind, which drastically lowers the probability to find them using paired recordings (conversely, in a different ongoing study we recorded closely connected FSI-SPN pairs with a high success rate–26 out 30, 87%). Therefore, our experiments do not directly answer the question whether a selected fraction *vs.* all of SST-INs are capable of co-releasing GABA and glutamate. In rtPCR analysis, co-expression of VGLUT1 and VGAT mRNA was found in 30% of our collected cytoplasmic samples, suggesting that co-transmission involves only a subset of SST-INs. However, although we performed control analyses that were useful to validate our results (e.g., all SST-INs expressing somatostatin, cortical principal cells expressing VGLUT but not VGAT, SPNs expressing VGAT but not VGLUT, and cholinergic interneurons expressing VGLUT3), our findings are based on a relatively small sample size, and it remains possible that some mRNA expression is under the detection threshold with our technique. It seems therefore premature to conclude that SST-INs should be subdivided into VGLUT^+^ *vs*. VGLUT^−^ subgroups. Thus, while providing an unequivocal evidence for the expression of VGLUT1 in some SST-INs, our molecular findings are to be as yet considered a proof of principle for what concerns a quantitative estimation.

### Physiological relevance and further considerations

What is the physiological meaning of simultaneous GABA and glutamate transmission? Concurrent expression of one major neurotransmitter and one or more neuromodulators (such as amines or peptides) is very common in the central nervous system, but co-transmission of GABA and glutamate–which normally exert opposite and counterbalanced effects through dedicated cell types–may seem puzzling at a first glance. However, the different kinetics of the two neurotransmitter described here (rapidly depressing AMPA-mediated EPSCs *vs*. persisting GABAergic IPSCs) are consistently reflected in the unambiguous firing response made of a brief excitation followed by a longer inhibition in postsynaptic cells. To some extent these properties recall the effects of classic feed-forward inhibition (FFI, typically exerted by dedicated interneurons and their GABAergic output shaping temporal dynamics of synaptic excitation provided by glutamatergic afferents (Buzsaki, 1984)), but using a different circuit configuration (Supplementary Figure S2). Striatal SST-INs appear to enclose both components of FFI thanks to the different short-term dynamics of EPSCs and IPSCs. The result is a sharp, temporally contained increase in postsynaptic firing frequency similar to classic FFI (Isaacson and Scanziani, 2011). Interestingly, these properties consolidate as the animal progresses into young adulthood, which may correlate with the acquisition of new skills, flexibility, or cognitive functions. The timing of SPN firing is associated with movement initiation, execution, or termination (Schultz and Romo, 1988; Cui et al., 2013; Jin et al., 2014), thus its fine regulation mediated by interneurons is of critical importance (Gernert et al., 2000; Gage et al., 2010). SST-INs and FSIs respond differently to cortical input due to dissimilarities in both pre-and postsynaptic elements (e.g., release probability, composition of AMPA receptors, presence *vs*. absence of NMDA receptors, and input resistance) (Gittis et al., 2010). This adds up to their different firing patterns, synaptic spatial distributions (perisomatic *vs*. dendritic contacts), and configuration of synaptic functions (GABA-mediated FFI *vs*. GABA/glutamate co-transmission). Altogether, these properties most likely provide a crucial contribution to the diversification and coordination of neuronal microcircuits and to a fine modulation of the striatal functional output.

An important issue that remains to be clarified is whether glutamate and GABA are released by the same or different vesicle pools, and whether these pools reside in the same or different presynaptic boutons (Zander et al., 2010; Tritsch et al., 2016). Although we did not investigate the microanatomy of these hybrid synapses, the differences in EPSC *vs*. IPSC short-term plasticity may in principle suggest the recruitment of separated pools of vesicles undergoing different release dynamics. It is alternatively possible, however, that the two neurotransmitters are co-packaged in the same vesicles, but in different quantities. A lower amount of glutamate molecules per vesicle would result in a quickly exhausted action at the postsynaptic side, though different availability/saturation of postsynaptic GABAergic and glutamatergic receptors and/or different neurotransmitter clearance from the synaptic cleft may also play a role. Further work employing electron microscopy and pharmacologic approaches will help answer these questions.

Finally, the interplay of GABA and glutamate transmission at co-releasing synapses may well be subject to a variety of physiological regulatory mechanisms in health –and anomalous functioning in disease. Interestingly, in pallido-habenular, co-releasing projections a shifted balance of the two neurotransmitters is associated with severely altered behavioral states such as depression (Shabel et al., 2014) and cocaine relapse (Meye et al., 2016). In the future it will be interesting to investigate if changes in the GABA/glutamate equilibrium at striatal SST-IN synapses similarly generate pathological abnormalities.

## Acknowledgements

Support for this work was provided by grants from the Telethon Foundation (GGP15096 to A.S. and GGP16234 to S.T.) and the Italian Ministry of Health (Young Investigator grant # GR-2013-02355540 to A.S.). F.B. is supported by the Jerome Lejeune Foundation and Fondazione Umberto Veronesi.

## Author Contributions

Conceptualization, S.C. and S.T.; Methodology, S.C., M.Z., M.R., F.B., A.S., and S.T.; Investigation, S.C., M.Z., M.R., F.B., A.S., and S.T.; Writing – Original Draft, S.T.; Writing – Review & Editing, S.C., M.Z., M.R., F.B., A.S., and S.T.; Funding Acquisition, A.S. and S.T.; Supervision, A.S. and S.T.

