## Supplemental Figures for "Somatostatin-Expressing Interneurons Co-Release GABA and Glutamate onto Different Postsynaptic Targets in the Striatum"

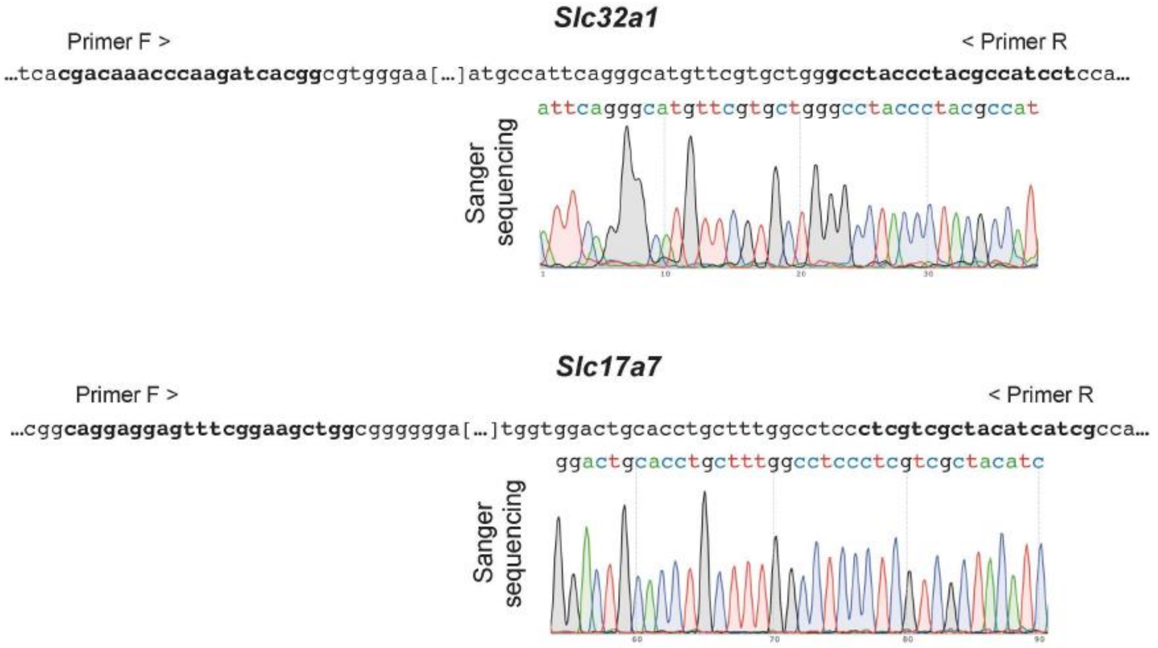


**Supplementary Figure S1. Sanger sequencing of single cell PCR.** Pherograms derived from Sanger sequencing of the PCR bands of *Slc32a1* and *Slc17a7*–amplified from the same SST-IN single-cell extract– exactly match with the relative gene coding sequences (primer sequences are shown in bold).

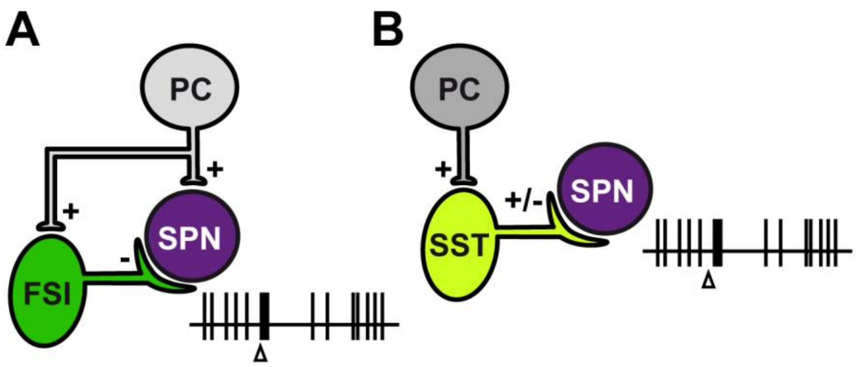


**Supplementary Figure S2. Feedforward inhibition and GABA/glutamate co-release may be segregated in differential cortico-striatal pathways and produce similar effects on SPN firing**. **A**: classic dysinaptic FFI configuration where a striatal SPN receives an excitatory input (+) from a cortical PC and an inhibitory one (-) from a neighbor FSI. An axon collateral from the PC also excites the FSI. The SPN responds to PC activation (arrowhead) with a transient increase in firing frequency–driven by monosynaptic excitation–followed by dysinaptic inhibition mediated by the FSI. **B**: in a different circuit, an SST-IN receives a glutamatergic input and projects a co-releasing synapse (+/-) to an SPN (not receiving a PC collateral), resulting in a similar firing response as that mediated by FFI (with a slight delay from PC activation due to the interposition of the SST-IN).
